# Systematic Conservation Planning for Intraspecific Genetic Diversity

**DOI:** 10.1101/105544

**Authors:** Ivan Paz-Vinas, Géraldine Loot, Virgilio Hermoso, Charlotte Veyssiere, Nicolas Poulet, Gaël Grenouillet, Simon Blanchet

**Affiliations:** Université de Toulouse, CNRS, UPS, IRD; UMR-5174 EDB, 118 route de Narbonne, 31062 Toulouse cedex 4, France; Aix-Marseille Université, CNRS, IRD, Avignon Université; UMR-7263 IMBE, 3 place Victor Hugo, 13331 Marseille cedex 3, France; Université de Lyon, CNRS, ENTPE; UMR-5023 LEHNA, 6 rue Raphaël Dubois, 69622 Villeurbanne, France; Institut Universitaire de France, Paris, France; Centre Tecnologic Forestal de Catalunya, Crta. Sant Llorenc de Monunys, Km 2, 25280, Solsona, Lleida, Spain; French Biodiversity Agency, pôle écohydraulique, Allée du professeur Camille Soula, 31400 Toulouse, France; CNRS, Station d’Écologie Théorique et Expérimentale; UMR-5321, 09200 Moulis, France

**Keywords:** biodiversity conservation, dendritic networks, multi-specific, microsatellites, conservation genetics, spatial biodiversity patterns

## Abstract

Intraspecific diversity informs the demographic and evolutionary histories of populations, and should be a main conservation target. Although approaches exist for identifying relevant biological conservation units, attempts to identify priority conservation areas for intraspecific diversity are scarce, especially within a multi-specific framework. We used neutral molecular data on six European freshwater fish species (*Squalius cephalus*, *Phoxinus phoxinus, Barbatula barbatula*, *Gobio occitaniae*, *Leuciscus burdigalensis* and *Parachondrostoma toxostoma*) sampled at the riverscape scale (i.e. the Garonne-Dordogne River basin, France) to determine hot- and cold-spots of genetic diversity, and to identify priority conservation areas using a systematic conservation planning approach. We demonstrate that systematic conservation planning is efficient for identifying priority areas representing a predefined part of the total genetic diversity of a whole landscape. With the exception of private allelic richness, classical genetic diversity indices (allelic richness, genetic uniqueness) were poor predictors for identifying priority areas. Moreover, we identified weak surrogacies among conservation solutions found for each species, implying that conservation solutions are highly species-specific. Nonetheless, we showed that priority areas identified using intraspecific genetic data from multiple species provide more effective conservation solutions than areas identified for single species or on the basis of traditional taxonomic criteria.

## Background

Biodiversity conservation is often addressed by identifying protected areas with high biodiversity and/or landscape values [1]. Conservation areas are generally identified as areas with high proportions of endemic, rare or iconic species [2]. Alternatively, conservation planning can be based on the concept of complementarity between conservation areas [3], and on cost-effectiveness analyses such as systematic conservation planning procedures (hereafter SCP [4]). SCP aims at identifying a number of complementary and *irreplaceable* sites best representing a predefined proportion of the biodiversity observed in a region and that should be managed for conservation in priority at a minimum cost.

There have been major conceptual and empirical leaps advocating the importance of considering species evolutionary history into conservation plans [5,6]. For instance, some authors included information on the phylogenetic history of species assemblages in SCP to preserve both species identities and their macro-evolutionary history [7–9]. However, genetic diversity observed at the *population level* (within species) has rarely been considered in SCP [but see 10–13]. *Intraspecific genetic diversity* (hereafter IGD) is a fundamental biodiversity facet, as it informs the demographic and colonisation history of populations (i.e. changes in effective population size over time and space), their level of biological connectivity, and their potential for adapting to environmental changes [14–16]. Conservation geneticists generally use neutral genetic diversity (i.e. genome portions unaffected by adaptive processes) to identify relevant evolutionary units within species [17], mainly by analysing genetic differentiation among populations [18,19]. For instance, the “Evolutionary Significant Units” (ESUs) framework has been proposed for identifying different lineages for conservation prioritization [20,21]. By emphasizing complementarity among areas –rather than differentiation–, SCP appears as a promising and complementary framework for identifying optimal sets of sites preserving pre-defined amounts of the total IGD observed within landscapes.

Most SCP studies yet to consider IGD aimed at identifying efficient surrogates for preserving genetic diversity [12,13]. For instance, two recent studies showed that protected areas identified using traditional species distribution data [13], and environmental and geographic descriptors [12] can be efficient surrogates in conservation planning for representing IGD. Although these studies shed light on how SCP can help preserve the IGD of multiple species, only a few studies have directly used multi-specific genotypic data for SCP (but see [11]). Datasets generated using neutral codominant genetic markers such as microsatellite markers are similar to datasets generally used for SCP (species presence/absence matrices), hence making possible applying SCP to genotypic surveys without needing intermediate modelling procedures or the calculation of summary genetic statistics (e.g. [10,12]) that may fail to capture the whole IGD of populations.

Here, we tested the potential of SCP to identify priority conservation areas accounting for IGD of a species assemblage at the landscape scale. We first considered a set of four common freshwater fish species (the chub *Squalius cephalus*, the gudgeon *Gobio occitaniae*, the stone loach *Barbatula barbatula* and the minnow *Phoxinus phoxinus*) to test the influence of conservation targets (proportion of the total amount of IGD to be secured by irreplaceable sites) and analytical strategies (analysing each species independently or all species pooled) on final conservation solutions (number and identity of irreplaceable sites). We then included two rare species of particular conservation interest (the beaked dace *Leuciscus burdigalensis* and the toxostome *Parachondrostoma toxostoma*) to test the relevance of the SCP approach in a “real conservation-oriented study”. For all species, we ran SCP analyses considering a typical conservation target [22] to explore the spatial distribution of irreplaceable sites in the riverscape, and to test for surrogacy in irreplaceable sites among species. We further tested whether irreplaceable sites identified using SCP are more efficient for preserving IGD than randomly-selected sites or than sites selected using traditional taxonomic criteria (e.g. sites displaying high species richness). We finally tested whether irreplaceable sites were correctly predicted by classical genetic diversity indices. We demonstrate that preserving the IGD of a species assemblage using systematic conservation planning is an efficient, yet complex, approach necessitating appropriate analyses to assist the decision-making process.

## Methods

### Biological models

We focused on six freshwater fish species: *Squalius cephalus*, *Phoxinus phoxinus* and *Barbatula barbatula,* which are widespread in Europe, and *Gobio occitaniae*, *Leuciscus burdigalensis* and *Parachondrostoma toxostoma*, which are endemic to Southern France [23]. These species were all from the Cyprinidae family apart from *B. barbatula,* which is a Nemacheilidae. This species assemblage covers a large functional trait space representative of many freshwater fish communities. For instance, *S. cephalus* is a large-bodied species with a long lifespan (i.e. up to 60 cm long and 15 years old [23]) whereas *Ph. phoxinus* is a small-bodied species with a shorter lifespan (i.e. less than 12 cm long and 4-5 years old [23]). From an ecological perspective, *G. occitaniae, Pa. toxostoma* and *B. barbatula* are bottom feeders, whereas *S. cephalus* and *Ph. phoxinus* are water column feeders. *Leuciscus burdigalensis* is more opportunistic. Further, *B. barbatula* is mainly active during the night, while the other species are active during the day. Four of these species are abundant (*S. cephalus*, *G. occitaniae*, *Ph. phoxinus* and *B. barbatula*), although their ecological niche and spatial occupancy in the Garonne-Dordogne River basin varies (see Figure S1). We will hereafter refer to this set of species as the “common” species. The two other species are rare (*L. burdigalensis*) to very rare (*Pa. toxostoma;* Figure S1) in the riverscape, and are of particular interest for conservation. *Leuciscus burdigalensis* is a recently described species that is experiencing both demographic and genetic bottlenecks in many populations [24,25]. *Parachondrostoma toxostoma* is a vulnerable species [26] listed in the IUCN red list, in the Annex II of the European Union Habitats Directive and in the Appendix III of the Bern Convention [26].

### Sampling design

During Spring/Summer 2010-2011, we used electrofishing to sample 92 sites distributed across 35 rivers from a whole European river basin (the Garonne-Dordogne River basin; >100,000 km^2^, South-Western France; Figure S1). Sampling sites were chosen to cover the entire distribution of each species in the riverscape, and to allow characterising their spatial patterns of IGD. Up to 25 individuals *per* species *per* site were sampled when possible. Not all species were present at all sampling sites (Figure S1; Table S1), and not all sites provided 25 individuals *per* species due to low densities. In these cases, we captured as many individuals as possible. We anesthetized each individual and then collected and stored in 90% ethanol a fragment of their pelvic fin. All individuals were released alive at their sampling location.

### Genotyping

Genomic DNA was extracted using a salt-extraction protocol [27]. We used multiplexed Polymerase Chain Reactions (PCRs) to co-amplify 8-15 microsatellite loci depending on the species (8 for *G. occitaniae*, 9 for *B. barbatula*, 10 for *S. cephalus* and *Ph. phoxinus*, 14 for *L. burdigalensis* and 15 for *Pa. toxostoma* (see Appendix A1 for details on recipes).

### Genetic diversity assessment

Given the good spatial resolution of the sampling obtained for the common species, descriptive genetic analyses were conducted for sampling sites displaying minimum sample sizes of N=10 individuals for these species to maximize consistency on subsequent allelic frequency-based genetic analyses. For the two rare species, for which the sampling was more restricted, genetic analyses were conducted for sampling sites displaying minimum sample sizes of N=6 individuals to maximize the number of sampling sites included in SCP. Reducing the minimum sample size from 10 to 6 permitted increasing considerably the number of sites retained for genetic analyses (21 to 29 sites, and 8 to 14 sites for *L. burdigalensis* and *Pa. toxostoma* respectively), hence improving the representation of their spatial distribution. We then determined for all six species the occurrence of null alleles, potential scoring errors, departures from Hardy-Weinberg equilibrium and linkage disequilibria among loci within sites (see Appendix A2 for details concerning these quality controls).

To assess within-site IGD (*α*-IGD), we applied the rarefaction procedures implemented in ADZE [28] to calculate both allelic richness (*AR*) and private allelic richness (*PA*) *per* sampling site (considering a minimum of N=10 individuals for common species or N=6 individuals for rare species). To assess among-site IGD (*β*-IGD), we used the R package ‘mmod’ [29] to calculate –for each species– a pairwise genetic differentiation index (*D*_*est*_ [30]). For each site (and species), we then derived the averaged value of all pairwise *D*_*est*_ values estimated between one given site and all the remaining sites to obtain a single value *per* site. For rare species, the impact of considering sites with N≥6 individuals instead of N≥10 on *AR*, *PA* and *D*_*est*_ was low (no clear impact neither on *PA* nor on *D*_*est*_, and mean decreases in *AR* of 0.295 and 1.183 alleles *per* site for *Pa. toxostoma* and *L. burdigalensis* respectively). This impact was however overcompensated by the increase in the number of sites retained for rare species when considering sites with N≥6 individuals.

#### Testing the suitability of SCP for IGD

In this first step, we tested the influence of conservation targets and analytical strategies on conservation solutions. We focused on data from the four common species, as their large coverage of the sampling area is more suited for the demonstrative exercise done in this step.

### Spatial patterns of IGD

We first used geostatistical modelling tools to explore spatial patterns of *α* -and *β*-IGD for the four common species in the riverscape by predicting *AR*, *PA* and *D*_*est*_ distributions from the observed empirical values using Generalized Linear Models for Spatial Stream Networks (GLMSSN [31,32]). These models were built by assuming three geographic descriptors (i.e. topological distance from the outlet, longitude and latitude) as explanatory variables (see Appendix A3 for further details). We used predictions from the best models (obtained through an Akaike Information Criteria model comparison) to produce kriged maps for each common species and each genetic index (Appendix A3). This allowed us to represent the spatial distribution of IGD across the whole river drainage, and hence to visually highlight hot- and cold-spots of IGD. Finally, we calculated Pearson’s correlation coefficients between *AR*, *PA* and *D*_*est*_ values calculated at the site level for each pair of common species to test for spatial congruence in IGD patterns among species.

### Identification of irreplaceable sites

We tested whether conservation targets (i.e. the percentage of total number of conservation units present in conservation solutions) and analytical approaches (i.e. species-specific or species-pooled analyses) influence the identification of irreplaceable sites. SCP methods traditionally use species presence/absence data as input data to identify irreplaceable sites (i.e. sites that cannot be excluded from an optimal selection of sites for conservation [33]) for the conservation of taxonomic diversity at the community level. Here, we replaced species presence/absence data with allele presence-absence data as in [11] to identify irreplaceable sites for the conservation of IGD of each species at the riverscape scale. We used Marxan v2.1 [33] and genotypic data from common species to identify, at the species level, an optimal set of sites best representing at least 50, 75, 90 or 100% of the total number of alleles present in the whole riverscape at a minimum “cost” (see Appendix A4 for further details). We then tested and compared visually how the proportions of irreplaceable sites vary among species and conservation targets.

To test how pooling data from several species affects the identification of irreplaceable sites, we further performed a “pooled” analysis, in which all alleles found for each common species at a site were pooled together in a single input dataset. We then selected random pulls of 30, 50, 75, 90 and 100% of the total number of alleles existing at the riverscape level (all common species confounded), and performed 100 Marxan runs *per* conservation target to identify the minimum set of sites representing these particular selections of alleles.

#### A real conservation-oriented study using SCP

In this second step, we (i) explored the spatial distribution of irreplaceable sites, (ii) tested whether conservation solutions found for each species are congruent among species, (iii) tested whether solutions found for one species can be used as a surrogate for other species, (iv) assessed the efficiency of the different conservation solutions we identified for preserving the overall IGD of all species, (v) compared the efficiency of conservation solutions identified through SCP to the efficiency of alternative conservation solutions based on traditional taxonomic criteria (e.g. in sites with high species richness), and finally (vi) tested whether irreplaceable sites are correctly predicted by classical genetic diversity indices. In this step, we included the two rare species, since surrogacy is particularly important to measure for rare species for which data are difficult to collect. We therefore focus more specifically on common *vs*. rare species comparisons.

### Identification of irreplaceable sites for genetic conservation

We here focused only on irreplaceable sites identified for the 90% target and used the program Marxan as described above to identify these sites for *L. burdigalensis* and *Pa. toxostoma* independently. We then mapped these irreplaceable sites on the river network to test (i) whether specific areas harboured more irreplaceable sites (e.g. upstream areas that are generally thought to be of high conservation priority [34]) and (ii) if irreplaceable sites are spatially congruent among species and, notably, among common and rare species. Additionally, we ran Generalized Linear Models (GLMs) including whether a sampling site is irreplaceable at the 90% target as a binomial dependent variable, and distance to the outlet of sites and *per* site betweenness centrality values [35] as explanatory variables. Betweenness centrality is an index quantifying the positional importance of each sampling site within the riverscape [36,37]. We assumed binomial error terms distributions and tested the significance of each term at the *α*=0.05 threshold.

### Surrogacy in irreplaceable sites among species

We estimated surrogacy levels among species by calculating the percentage of the total number of alleles observed for a given species secured by irreplaceable sites identified for another species. Although surrogacy was calculated for all species pairs, we specifically focused on rare species by calculating (i) the percentage of the total number of alleles observed for rare species secured by irreplaceable sites identified for all common species, and (ii) the percentage of the total number of alleles observed for common species secured by irreplaceable sites identified for all rare species.

### Assessing the efficiency of SPC

The goal of this analysis was (i) to assess the efficiency of SCP for preserving the overall levels of IGD observed in the riverscape for the six species, and to (ii) compare the efficiency of SCP-based conservation solutions to the efficiency of alternative conservation solutions based on traditional taxonomic criteria. We first computed an average accumulation curve of “protected genetic diversity” by estimating the average cumulative percentage of the total number of alleles (over all species) secured by a cumulative number of randomly-selected sampling sites. Specifically, we increased the number of sampled sites from 1 to 92 with a continuous increment of one unit. At each incremental step, we randomly selected sites (without replacement) and calculated the percentage of the total number of alleles (over all species) secured by the random set of sites. We repeated this random selection 1000 times and averaged the percentages over these 1000 permutations to obtain the averaged accumulation curve of protected genetic diversity. We then tested to which extent, for an identical number of sites, the conservation solutions we identified through SCP (i.e. irreplaceable sites) secured higher percentages of IGD than those secured by a random selection of sites. Notably, we explored whether certain conservation solutions (i.e. by considering irreplaceable sites identified for single species or for the sets of common, rare and all six species) were more efficient than others to sustain a large percentage of IGD.

As in most conservation studies species diversity is the main conservation target, we finally checked the efficiency of conservation solutions based on taxonomic criteria for preserving the overall IGD of the six studied species. Specifically, we calculated the percentages of total number of alleles (over all species) secured by sites (i) in which a number equal or greater than 10, 15 or 20 freshwater fish species co-occur (the total number of fish species observed in the riverscape being 34, with a mean number of species observed at the site level of 12.77 ±0.55), and (ii) in which the six studied species co-occur. We finally compared the efficiencies of taxonomic-data-based and SCP-based conservation solutions to assess if solutions based on taxonomic criteria are good surrogates of the overall IGD of all species. We derived data on fish species occurrence from [37] for 81 of our 92 sampling sites.

### Relationships between irreplaceable sites and genetic diversity indices

To test the ability of classical genetic indices to predict the propensity of sites to be irreplaceable, we ran for each species GLMs (considering binomial error terms distributions) including whether sites were designated as irreplaceable at the 90% target as a binomial dependent variable, and *AR*, *PA* and *D*_*est*_ as explanatory variables. We tested the significance of each term at the *α*=0.05 threshold. Explanatory variables were centred and scaled to compare the relative strength of the predictors among species.

## Results

### Descriptive statistics

Overall, *Pa. toxostoma* showed the lowest within-site IGD estimates. Mean *AR* ranged from 2.114 for *Pa. toxostoma* to 5.821 for *Ph. phoxinus*, and mean *PA* ranged from 0.036 for *Pa. toxostoma* to 0.162 for *L. burdigalensis* (Table S2). *Parachondrostoma toxostoma* also showed the lowest mean *D*_*est*_ value (0.069), while *B. barbatula* displayed the highest mean value (0.383; Table S2).

#### Testing the suitability of SCP for IGD

##### Spatial patterns of IGD

Spatial patterns of IGD largely varied and were poorly congruent among common species (Figure 1). For instance, hotspots of *AR* for *S. cephalus* were mainly found in the Western part of the riverscape and on core streams, whereas these same areas were identified as coldspots of *AR* for *B. barbatula* (Figure 1A1-1A3). Similarly, *PA* hotspots were inversely related between *G. occitaniae* and *Ph. phoxinus* (Figure 1B2-1B4). Similar conclusions were reached for *D*_*est*_ (Figure 1C). For instance, *D*_*est*_ hotspots were observed in opposite riverscape areas for the species pair *B. barbatula/Ph. phoxinus* (Figure 1C3-1C4). Consequently, the sign, slope and significance of GLMSSNs explanatory variables strongly varied among species (Figure 1). This incongruence in spatial patterns of IGD among species was also reflected by the low to moderate correlation coefficients measured among all possible species pairs and for each genetic diversity index (i.e. *AR*, *PA* and *D*_*est*_, Table S3). Indeed, Pearson’s correlation coefficients were lower than 0.6 for all comparisons but two (i.e. between *B. barbatula*/*G. occitaniae* and between *Ph. phoxinus*/*B. barbatula* for *AR*, Table S3).

**Figure 1.**
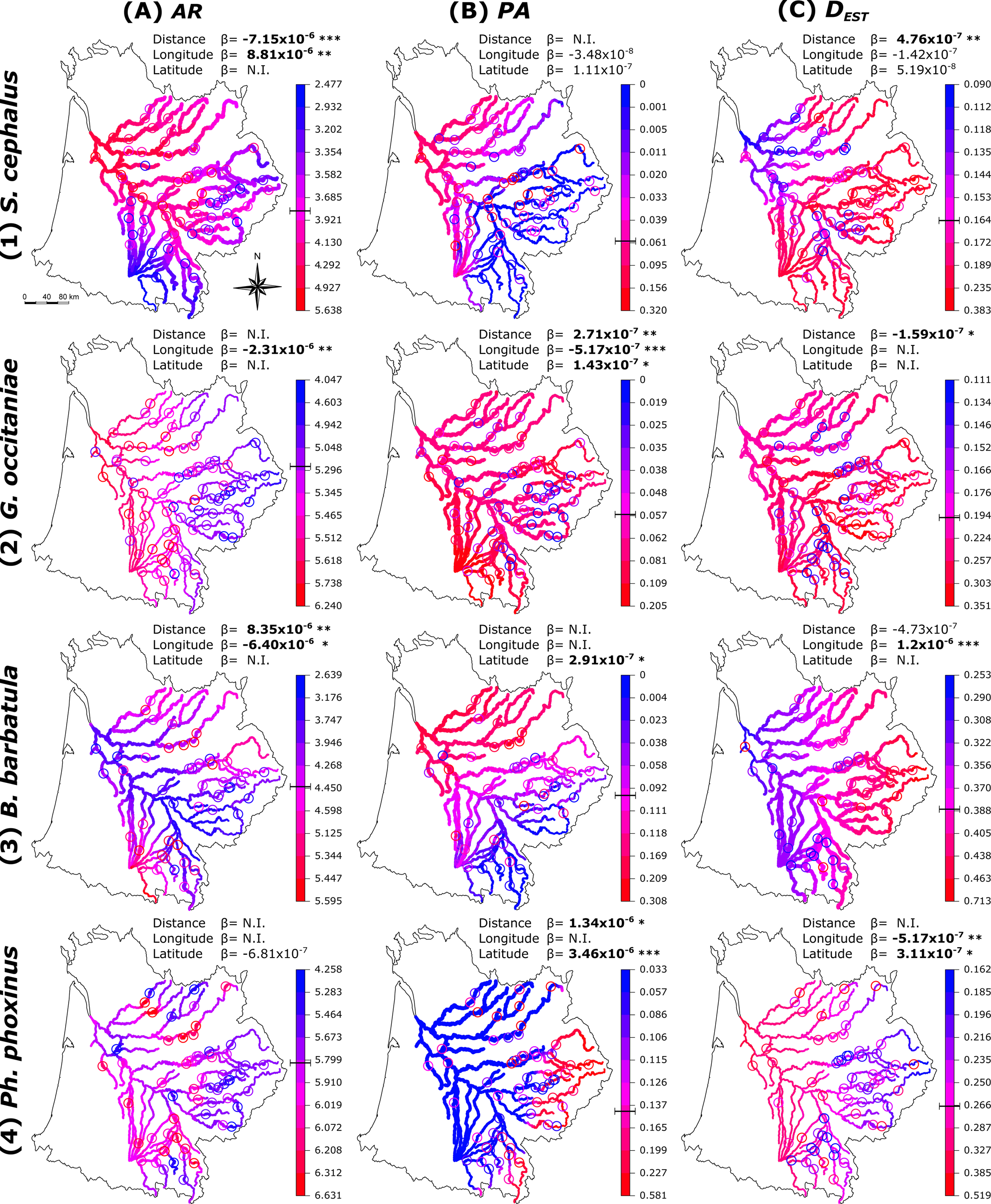
Spatial distribution of observed (coloured circles) and interpolated (coloured lines) values of *AR*, *PA* and *D*_*est*_ (A, B, C respectively in the figure) for *Squalius cephalus*, *Gobio occitaniae*, *Barbatula barbatula,* and *Phoxinus phoxinus* (1, 2, 3, 4 respectively in the figure) obtained with GLMSSNs. Coloured lines width is inversely related to prediction standard error. The cursor on the vertical coloured scale indicates mean *AR*, *PA* and *D*_*est*_ values. The slope (β) of each explanatory variable (topological distance to the outlet, longitude and latitude) and its significance is also reported. N.I. indicates that the explanatory variable was not included in the model; *p-value<0.05; **p-value<0.01; ***p-value<0.001.

##### The influence of conservation targets and analytical strategies to identify irreplaceable sites for genetic conservation

When species were analysed independently, we found that the number of irreplaceable sites increased as the conservation target increased, with a steep increase for conservation targets higher than 75% of the total number of alleles (Figure 2). However, the percentage of irreplaceable sites strongly varied among species. For instance, for a 90% target, the proportion of irreplaceable sites ranged from 3.61% of the total number of sampled sites for *G. occitaniae* to 28.57% for *Ph. phoxinus* (Table S4; Figure 2). For extreme conservation targets (100% of alleles to be secured), the proportion of irreplaceable sites varied from 25.30% for *G. occitaniae* to 68.26% for *Ph. phoxinus* (Table S4; Figure 2).

**Figure 2.**
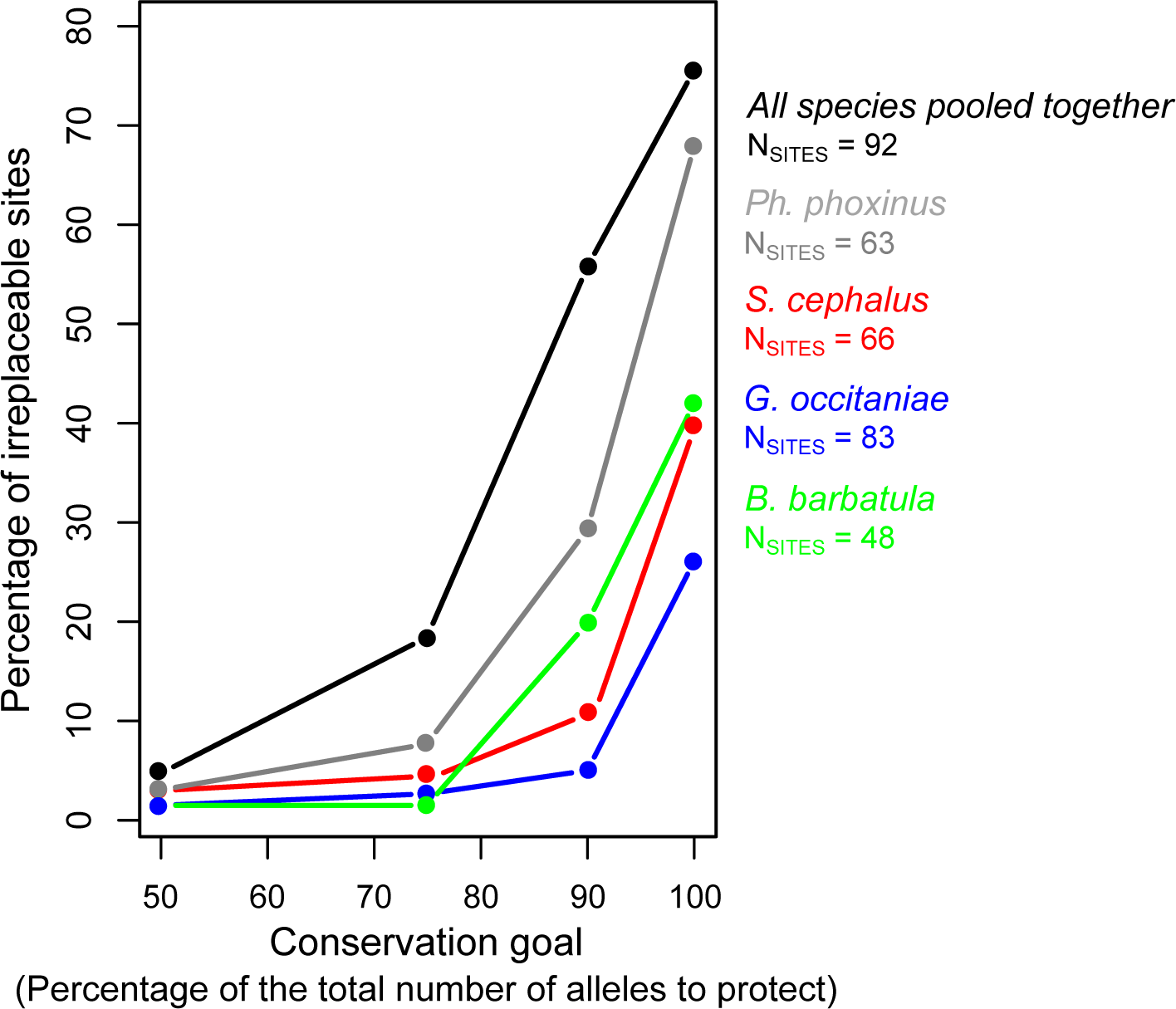
Percentage of irreplaceable sites identified by Marxan for conservation targets of 50, 75, 90 and 100% of the total number of alleles present in the riverscape for each common species and for a pooled analysis in which all alleles from all common species were pooled together. N_SITES_ represents the number of sites included in Marxan analyses.

When alleles from the four common species were analysed in a single pooled analysis, we similarly found that the proportion of irreplaceable sites increases as the conservation target increases (Figure 2). Interestingly, no irreplaceable sites were identified for the 30% target, and only 3 irreplaceable sites were found for the 50% target (Figure 2). The proportion of irreplaceable sites increased moderately to 17.39% for the 75% target, and then steeply increased for higher conservation targets to reach 55.70% for the 90% target and 76.08% for the 100% target (Figure 2). This later result suggests that almost all the riverscape should be protected to reach high conservation targets when adopting a pooled strategy.

#### A real conservation-oriented study using SCP

##### Identification of irreplaceable sites for genetic conservation

We first visually explored the spatial distribution of irreplaceable sites. Overall, the localization of irreplaceable sites in the riverscape strongly varied among species (Figure S2). We failed to identify areas (e.g. upstream/downstream locations) clustering irreplaceable sites for any species (Figure S2A-F). This apparent lack of clustering was statistically confirmed by GLMs (Table S5). Indeed, neither the distance from the outlet of sampling sites nor their betweenness centralities were significant predictors of sites irreplaceability for all species (excepting a significant effect of distance from the outlet for *Ph. phoxinus*, Table S5).

Second, we tested whether conservation solutions found for each species were spatially congruent among species. Over the six fish species, forty-two sites (45.65%) were irreplaceable at the 90% conservation target for at least one species (Figure 3). Thirty-two of these forty-two sites were irreplaceable for at least one common species (Figure 3), and fourteen of the forty-two sites were irreplaceable for at least one rare species (Figure 3). Among the six species, only eight of these forty-two sites were irreplaceable for at least two species, and only one site was irreplaceable for three species (Figure 3).

**Figure 3.**
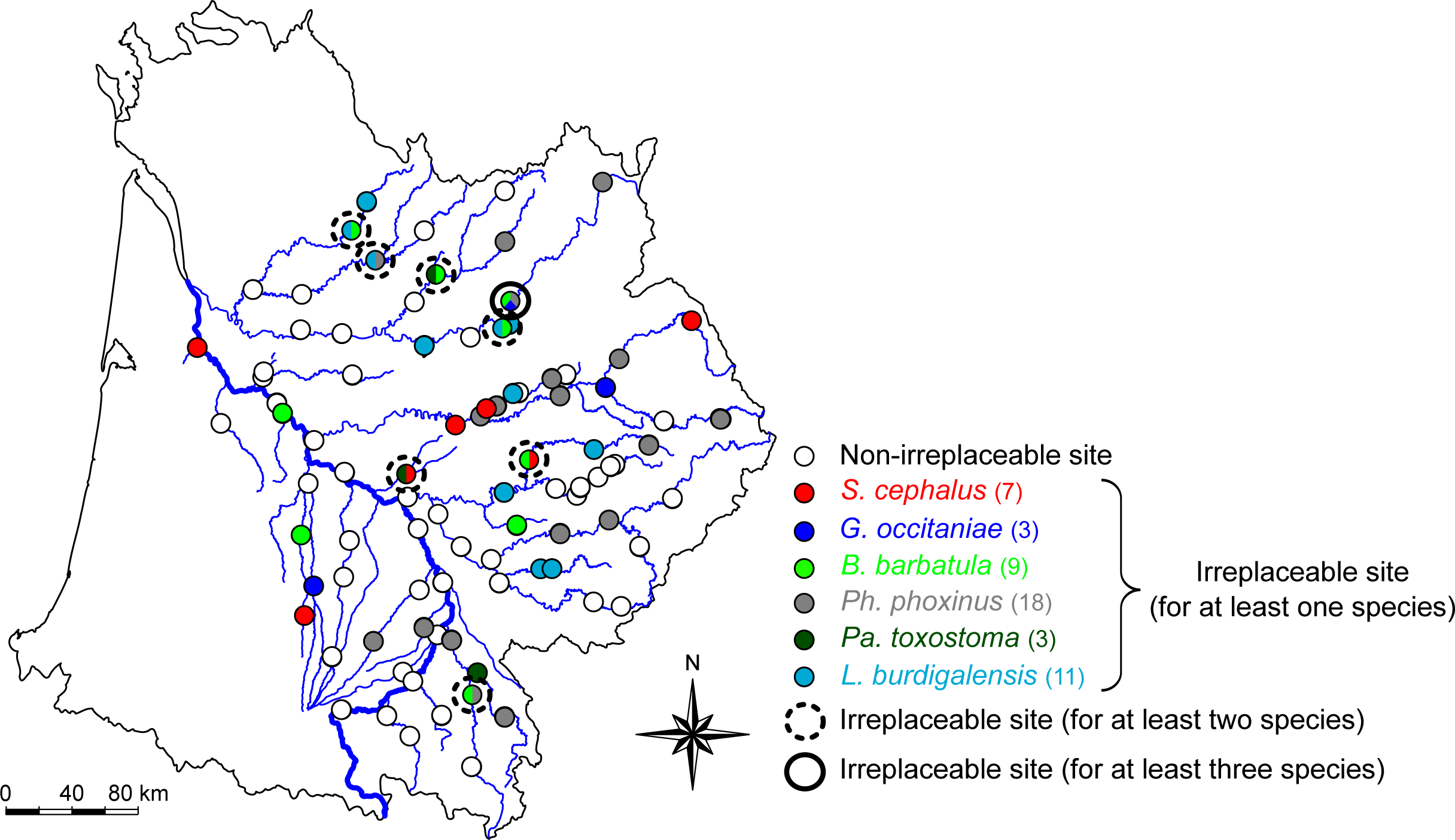
Irreplaceable sites identified by Marxan for at least one (unicoloured-filled points), two (black dotted circles surrounding bicoloured points) or three (black bolded circle surrounding a tricoloured point) species, assuming a conservation target of 90% of the total number of alleles present in the riverscape.

##### Surrogacy in irreplaceable sites among species

Surrogacy levels were generally low to moderate, and strongly varied among species pairs (Table S6). For instance, irreplaceable sites identified for *G. occitaniae* failed to secure rare species IGD, and covered only 18 to 43% of the total number of alleles of the other common species. This indicates that irreplaceable sites identified for *G. occitaniae* (the most widespread species) are poor surrogates for preserving the IGD of other species (Table S6). Conversely, irreplaceable sites found for *Ph. phoxinus* and *L. burdigalensis* are better surrogates for *G. occitaniae*, as 79.58% and 70.27% of the total number of alleles of *G. occitaniae* was secured by irreplaceable sites identified for *Ph. phoxinus* and *L. burdigalensis* respectively (Table S6). Overall, irreplaceable sites best sustaining other species IGD were those identified for *B. barbatula*, which secured in average 68.28% of the total number of alleles of other species (Table S6).

Interestingly, the thirty-two irreplaceable sites identified for common species secured 81.79% and 90% of the total number of alleles of *L. burdigalensis* and *Pa. toxostoma* respectively, suggesting that irreplaceable sites identified for a set of common species can be good surrogates for rare species IGD. The fourteen irreplaceable sites identified for the two rare species secured a total number of alleles ranging from 65.18% for *S. cephalus* to 79.40% for *B. barbatula*, which is not negligible given that conservation priorities are generally directed towards rare species.

##### Assessing the efficiency of SPC

Overall, irreplaceable sites identified for single species using SCP did not significantly secured higher percentages of IGD than randomly-selected sites, excepting irreplaceable sites identified for *L. burdigalensis* and *B. barbatula* (Figure 4), which were more efficient than randomly-selected sites for preserving the overall IGD of all species. However, we found significantly higher percentages of secured alleles (compared to randomly-selected sites) on irreplaceable sites identified through SCP for (i) the set of common species (ii) the set of rare species, and (iii) all the six species (Figure 4).

**Figure 4.**
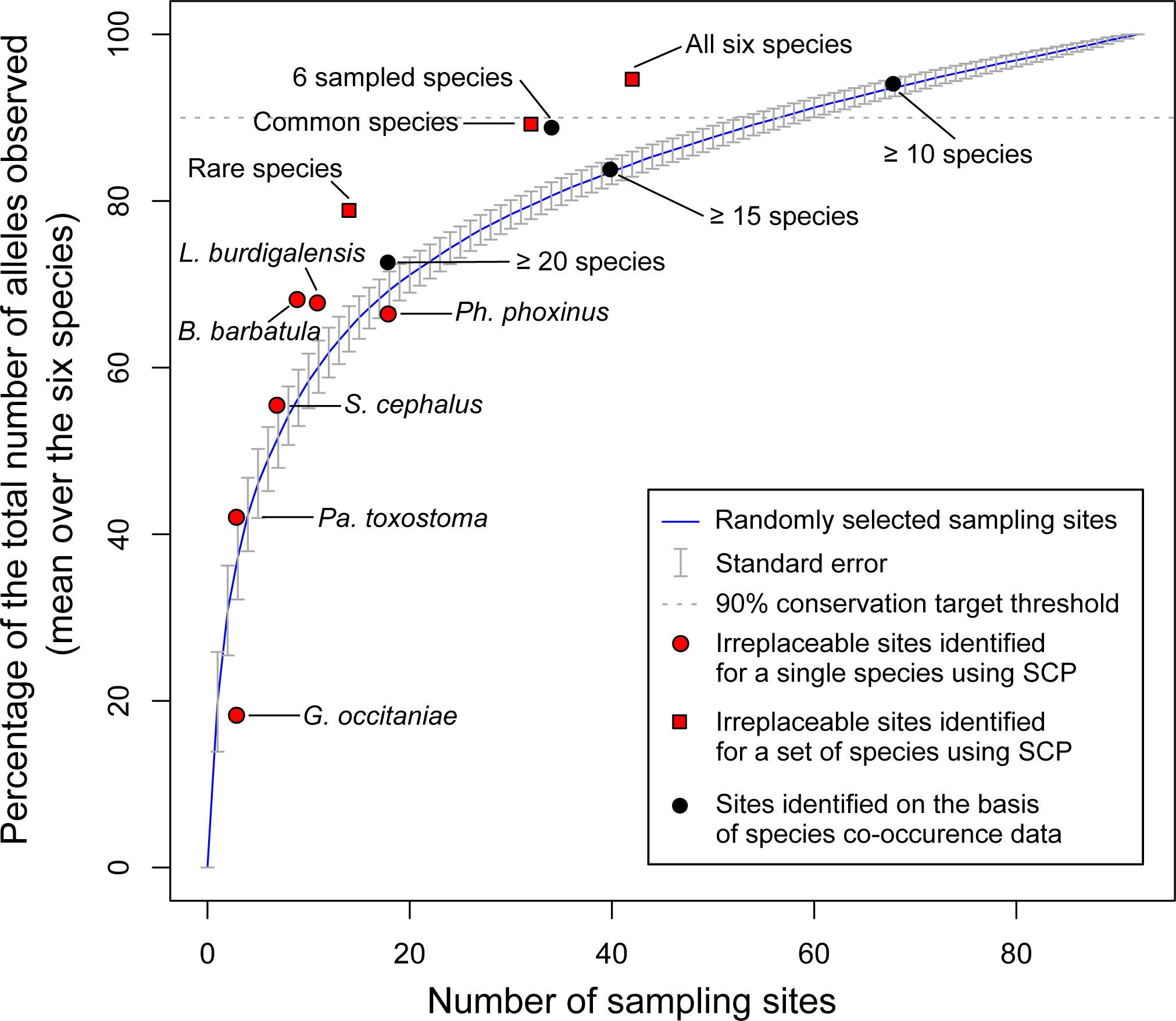
Mean percentage of the total number of alleles observed in the riverscape for the six species that are secured by (i) a cumulative number of randomly-selected sampling sites (blue line; average accumulation curve obtained under 1000 permutations of the sampling sites; grey vertical lines: standard error), (ii) irreplaceable sites identified using SCP for each species independently (red dots), (iii) irreplaceable sites identified using SCP for different sets of species (red squares), and (iv) sites identified on the basis of freshwater fish species co-occurrence data (black dots).

Conservation solutions based on the co-occurrence of high numbers of species (i.e. sites with numbers of co-occurring species equal or greater than 10, 15 or 20) did not secured more alleles than randomly-selected sites, suggesting that raw interspecific diversity (i.e. species richness) may not be a good surrogate of IGD in our study (Figure 4). Nevertheless, we found that a higher percentage of alleles would be secured by selecting sites on which the six species co-occur (compared to randomly-selected sites; Figure 4).

##### Relationships between genetic indices and irreplaceable sites

For all species but *G. occitaniae* and *Pa. toxostoma* (for which no variables were significant predictors), *PA* was identified as the only variable significantly predicting the probability of sites to be irreplaceable (Table S7). This probability increased with the number of *PA* in a site.

## Discussion

Intraspecific genetic diversity reflects the evolutionary and demographic history of populations, and hence sustains the capacity of species and populations to cope with environmental changes [14–16]. This biodiversity facet is the first that should respond to global change [15], and it is hence surprising that IGD has yet been rarely integrated in dedicated optimization planning tools. We here fill this gap by demonstrating that systematic conservation planning for IGD is an efficient, yet complex, approach necessitating careful considerations.

### From idiosyncratic IGD distributions…

Our results suggest that the IGD of the targeted species did not follow a common spatial pattern, but rather species-specific (idiosyncratic) spatial distributions. This conclusion holds true for all genetic diversity indices, and corroborates the few previous studies investigating simultaneously the spatial distribution of IGD at large spatial scales and for sympatric species [38,39]. This was however unexpected given that recent meta-analyses on freshwater organisms demonstrated that *α*-IGD is generally higher in downstream than in upstream areas [40], and that *β*-IGD tends to be higher in upstream than in downstream sections [34]. Overall, these general spatial patterns of IGD were verified in our datasets. For instance, a negative relationship between allelic richness and distance from the outlet is expected in freshwater organisms [40], and was actually observed for three of the six studied species (Figure S3). However, when using a precise and novel approach to map genetic diversity across the network, we demonstrated that the distribution of cold- and hot-spots of *α*-and *β*-IGD was subtler and idiosyncratic. This probably reflects interactions between recent colonization histories, species life-history-traits and the network structure, which are expected to drive neutral IGD patterns in rivers [40–42].

### …to systematic conservation planning for IGD

The spatial mismatch in IGD among species probably explains why we found that the surrogacy level among irreplaceable sites identified for each species was extremely low. For instance, an extremely low proportion of irreplaceable sites were common to two or three species (and never more than three species; Figure 4). In the same way, we detected no clear patterns in the spatial distribution of irreplaceable sites. Indeed, irreplaceable sites (for any of the six species) were not particularly situated in specific locations such as upstream areas and/or in areas of high connectivity (i.e. with high centrality values [36,37]), hence suggesting that priority areas for the conservation of multi-specific IGD should not be restricted to specific areas in the riverscape. It is however noteworthy that the greatly variable position of irreplaceable sites found for the studied species may reflect both their spatial occupancy in the riverscape and their own ecological niche. In this demonstrative study, we only focused on the influence of spatial variables (longitude, latitude, distance to the outlet, centrality) on conservation solutions. Considering other environmental variables that may directly or indirectly influence the distribution of IGD such as water temperature, oxygen concentration and/or water pollution (e.g. [43]) would certainly shed important light on the drivers of sites irreplaceability, and should hence be the focus of future SCP studies. Finally, the surrogacy level among irreplaceable sites was low to moderate, and never attained the 90% threshold we assumed when we considered all species independently. Combined with our finding that the number of irreplaceable sites can be high for reasonable conservation targets (up to 46% sites identified as irreplaceable for at least one species at the 90% conservation target), we conclude that our ability to identify priority areas for IGD may be highly species-specific and may depend on the capacity to tackle the trade-off between the amount of IGD to protect, and the extent of priority areas we can realistically protect.

However, when surrogacy between all irreplaceable sites identified for the entire set of common species and those identified for the rare species was tested, we reached reasonable proportions of the total number of alleles to protect (~80-90%) for the two rare species (i.e. *Pa. toxostoma* and *L. burdigalensis*). This result suggests that, in some cases, genetic data obtained for a set of widely-distributed, “easier-to-sample” common species displaying varying life-history traits can be used for identifying protection areas for the IGD of other rare species that can be more problematic to sample. This would require, however, a good overlap between the spatial distributions of both the common species used as surrogates and the targeted rare species, and a wide sampling design allowing representing the whole spatial occupancy of species in the landscape. Failing to fulfil these two prerequisites may have a negative impact on conservation costs, as the number of irreplaceable sites necessary to preserve a predefined amount of IGD of rare species would be higher than expected. In our demonstrative study for instance, thirty-four of the forty-two (81%) irreplaceable sites were sites in which at least four study species co-occur, and nineteen of these sites (54.3%) were sites in which the six species co-occur. It is worth mentioning that our study builds on a large-scale (but scattered) sampling design to test SCP for IGD at the riverscape scale. The efficiency of SCP for preserving IGD at finer sampling scales (e.g. by concentrating sampling sites within a restricted landscape area) remains however to be tested. Another important finding we report here is that selecting conservation areas based on traditional taxonomic criteria (i.e. by selecting sites with high species richness) is not an efficient strategy for preserving the overall IGD of a set of species. This highlights the need for finding efficient ways for considering IGD in SCP, as we and others [9–13] proposed. Here, we have focused on six freshwater fish species, among which five were cyprinids. Future works should consider wider taxonomic diversities (e.g. by also considering IGD from vertebrate, invertebrate and/or plant species in SCP) to explore the distribution and drivers of site irreplaceability at a large taxonomic scale.

Overall, our results suggest that two different analytical strategies can be employed in real-case SCP studies aiming at preserving IGD of a species assemblage: (i) identifying conservation areas for a set of rare species or (ii) identifying conservation areas for a set of representative common species. Both strategies have their own advantages and inconveniences. The first strategy may optimally preserve IGD of rare species at competitive costs (e.g. only 14 irreplaceable sites to protect in our study), but this at the expense of the IGD of other sympatric species. Conversely, the second strategy may optimally preserve IGD of a set of common species while maintaining high levels of IGD for rare species, but this at a higher cost (32 irreplaceable sites to protect in our study). Whether to choose one of these two strategies will therefore depend on many factors such as how difficult is the sampling of rare species compared to common species, or the extent of the resources available for setting new protected areas. We recommend adopting the second strategy when possible, since it allows simultaneously maintaining IGD from rare and common species. Indeed, IGD of common species is vital for ensuring ecosystem stability, as it ultimately influences species interactions, population dynamics and ecosystem functioning [44]. It is noteworthy that further studies must be conducted to test if the two analytical strategies we propose can be applicable across taxa and landscapes.

Although it was not a main goal of the study, we identified an alternative strategy that may also be efficient for maintaining high levels of IGD for a species assemblage without requiring SCP analyses. Indeed, selecting sites where all the studied species co-occur as conservation solutions allowed maintaining high levels of IGD for both common and rare species in our study, although this strategy was less efficient than selecting irreplaceable sites found for common species through SCP. The efficiency of this strategy also needs to be thoroughly tested across taxa and landscapes, as it can be a useful surrogate strategy in some cases (e.g. when molecular data cannot be produced due to funding or technical limitations). It is however noteworthy that such strategy requires highly reliable taxonomic data gathered over the whole studied landscape.

We also demonstrate that microsatellite-based genotypic data can be easily used as direct input data in SCP to identify efficient conservation areas for IGD of a species assemblage at the landscape scale. Relatively small numbers of microsatellite markers such as those considered in our study (8-15) have been proven to be sufficient for capturing the confounding genetic effects of recent past demographic history, genetic drift and connectivity issues between populations [25,40,41], and their efficiency for informing recent evolutionary processes can be equal or higher than low to moderate numbers of SNP markers for instance (e.g. <300 SNPs; [18]). However, microsatellite markers have the disadvantage of being unsuitable for drawing conclusions on populations’ adaptive potential, contrarily to other markers derived at the genome scale. Further, the sample size threshold we set (up to 25 individuals *per* site and species) was sufficient to properly estimate IGD of the species assemblage at the riverscape scale without implying an unreasonable sampling effort. We recall, however, that genetic diversity was probably slightly underestimated for sites where small numbers of individuals were captured (e.g. sites with N=6 individuals for rare species; see the Methods section for details). Since our capacity to compile genetic datasets at large spatial, temporal and taxonomic scales greatly increased in recent decades, a next methodological step may be using high-throughput techniques to develop high numbers of markers informing both neutral and non-neutral processes to assess whether SCP solutions obtained using different markers allow efficiently preserving the whole genomic diversity of populations, and hence their whole evolutionary history.

### Conclusions

Our study provides novel, insightful and promising knowledge on the setting of priority conservation areas for intraspecific diversity. It shows that systematic conservation planning methods are useful and efficient objective tools for conservation geneticists whose conservation solutions will strikingly depend on the species to be conserved and the quantity of genetic information that managers aim at preserving in a landscape. Given our results, we suggest that two strategies could be employed in real-case conservation programs: (i) identifying priority conservation areas for a set of rare species when resources allocated to conservation are scarce or (ii) identifying priority conservation areas based on the analysis of a set of representative common species that may serve as “umbrellas” for rare sympatric species when resources allocated to conservation are higher.

Our study also raises many additional questions that should be considered in the near future. We believe that the next steps will be to formally identify sound conservation targets for intraspecific diversity, to test whether neutral intraspecific diversity appropriately mirrors quantitative and adaptive diversity [16], and to quantify the influence of intraspecific diversity on ecosystem functioning and services to better evaluate the added value of preserving this biodiversity facet [16,44].

## Ethics statement

Sampling was made in accordance with current French ethical laws (permit numbers 2010-161-10, 2010-108, 2011-185-06 and 2011-318).

## Data accessibility

Supplementary tables, figures and appendices are provided as Electronic supplementary material. We also provide raw allele counts and genetic diversity estimates for all species, and raw outputs for Hardy-Weinberg equilibrium, linkage disequilibrium and null allele’s tests.

## Authors’ Contributions

SB, GL and IPV designed the study. IPV and SB wrote the manuscript with the help of VH, GG, NP and GL. IPV, SB, GL and CV collected the samples. GL and CV produced the genetic data. IPV, VH and SB conducted statistical and population genetics analyses.

## Acknowledgements

We thank all our colleagues and ONEMA staff that contributed to the field sampling, especially Laurence Blanc. Lisa Fourtune is acknowledged for having calculated betweenness centrality values. We also thank Marie-Hélène Lizée, Camille Pagès and Jérôme Prunier for their comments, the “Génopole Toulouse” for help with genotyping and Keoni Saint-Pée for correcting the English. We are grateful to Loeske Kruuk, Robert Wilson and five anonymous reviewers for their comments on earlier versions of the manuscript. This work was supported by the “Laboratoire d’Excellence” (LABEX) entitled TULIP (ANR-10-LABX-41).

