## Supplementary Materials for "Systematic Conservation Planning for Intraspecific Genetic Diversity" (from another version of this preprint: version 4, 2018-03-17)

**FIGURE S1:** Maps of the Garonne-Dordogne river basin (South-Western France) representing the spatial distribution of the sampling sites where *Squalius cephalus* (A), *Gobio occitaniae* (B), *Barbatula barbatula* (C), *Phoxinus phoxinus* (D), *Leuciscus burdigalensis* (E) and *Parachondrostoma toxostoma* (F) have been successfully (green circles) or unsuccessfully (black circles) sampled for genetic analyses. Sites successfully sampled for genetic analyses are sites where at least ten individuals (or six individuals for the two rare species *L. burdigalensis* and *P. toxostoma*) have been successfully genotyped. These spatial distributions of the sampling sites roughly correspond to the spatial distribution of these species in the riverscape.

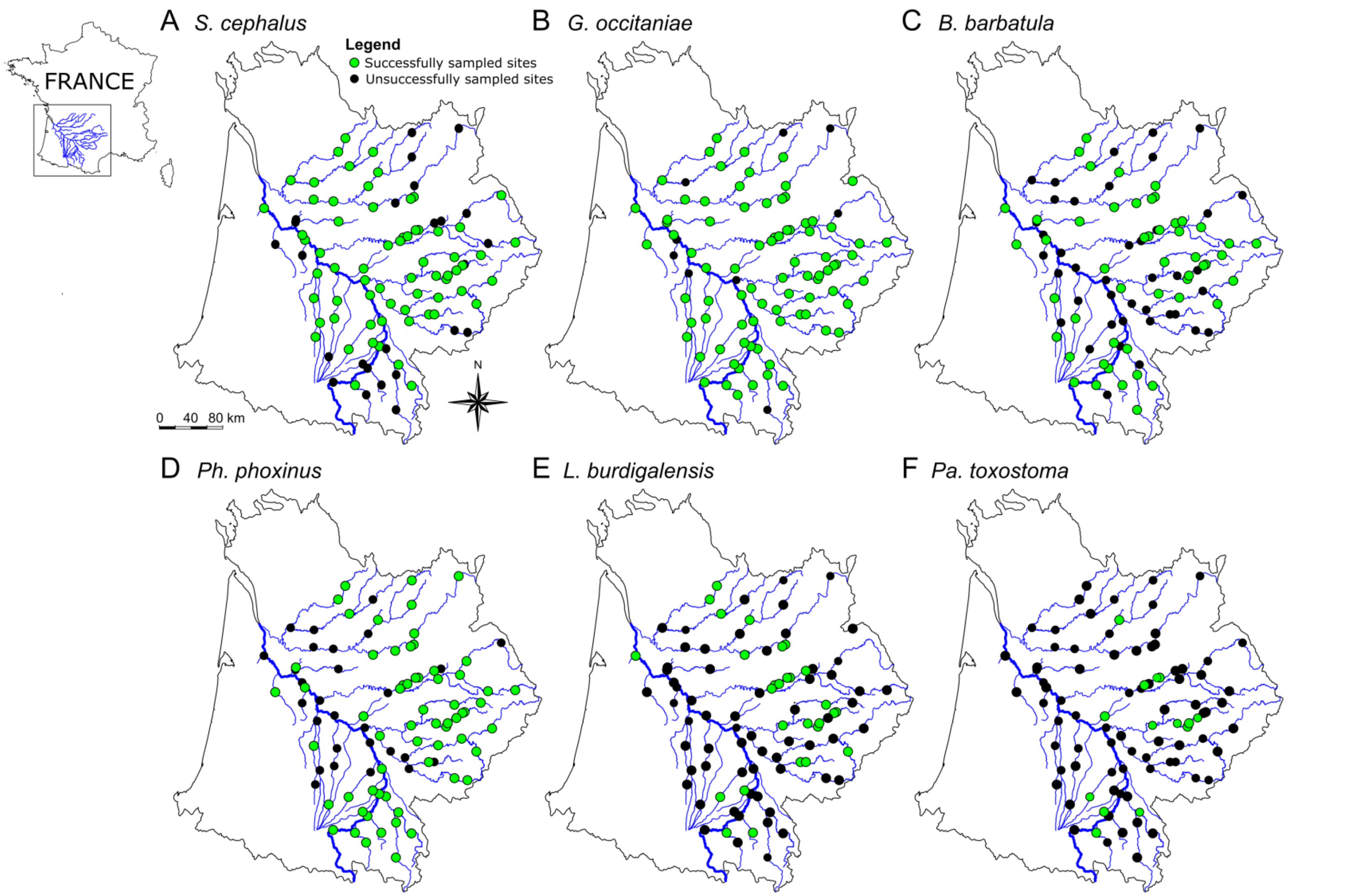

**FIGURE S2:** Irreplaceable sites that have been identified for each species by Marxan for preserving 90% of the total number of alleles present in the river basin (red-filled circles).

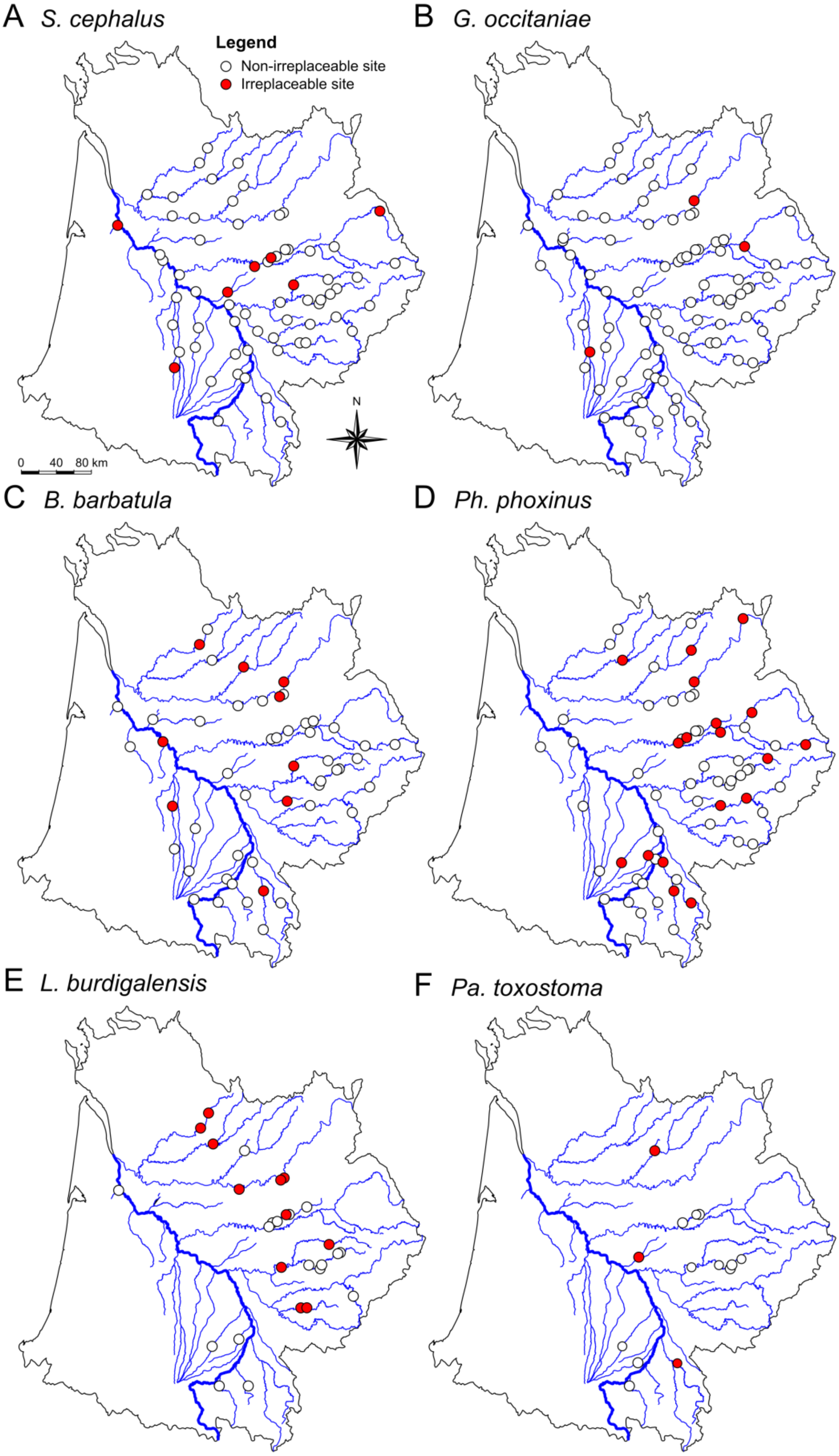

**FIGURE S3:** Allelic Richness (*AR*) *per* sampling site in function of the topological distance from the outlet of sampling sites (in km) for (A) *Squalius cephalus*, (B) *Gobio occitaniae*, (C) *Barbatula barbatula*, (D) *Phoxinus phoxinus*, (E) *Leuciscus burdigalensis* and (F) *Parachondrostoma toxostoma*. The black line corresponds to the predicted values obtained from linear models. In these models, *AR* was the dependent variable, and the topological distance from the outlet was the explanatory variable. The *r* values correspond to Pearson's correlation coefficients between the two variables. Significant values are in bold. ***** indicates p-values <0.05; ****** indicates p-values <0.01; ******* indicates p-values <0.001.

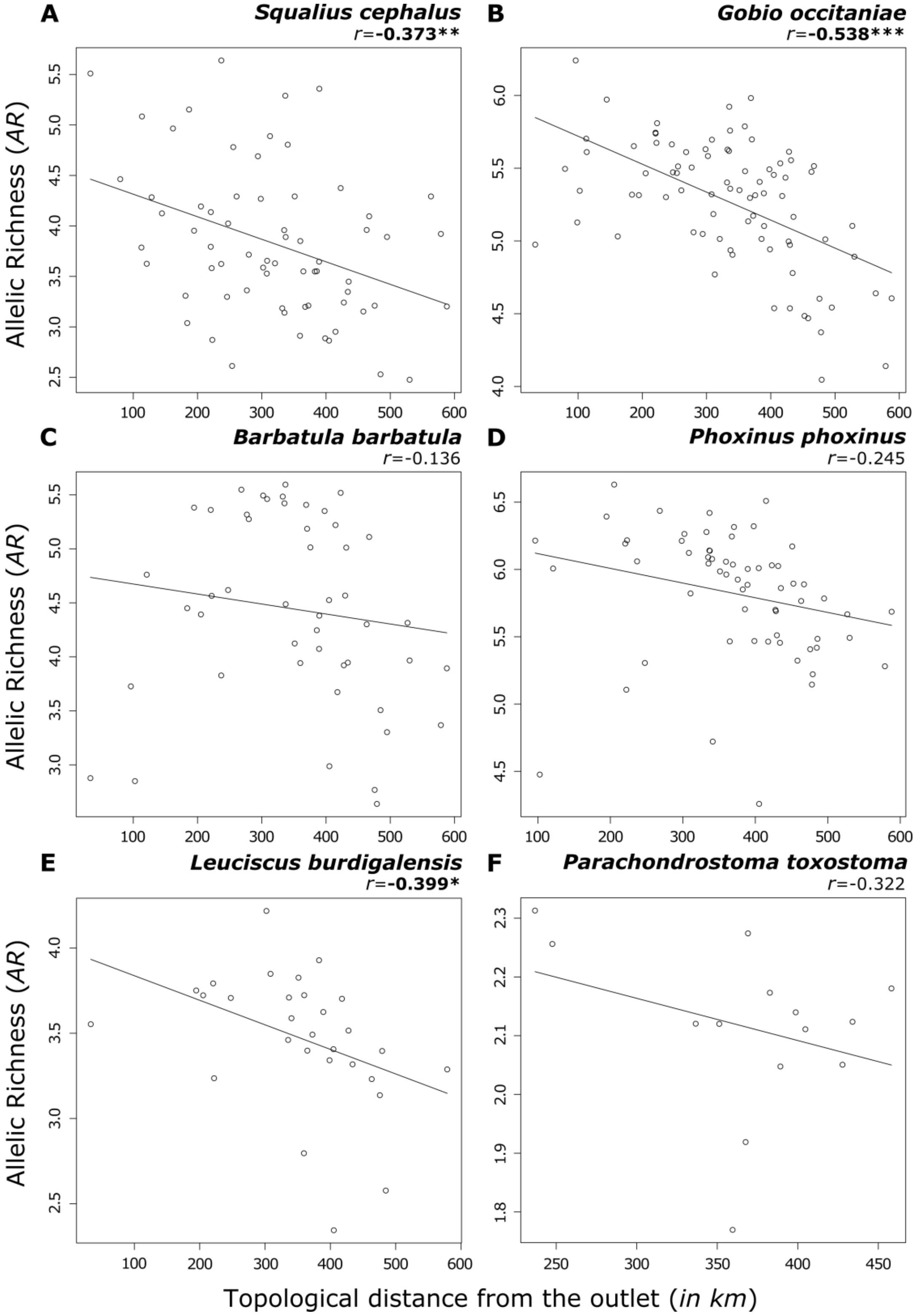

**TABLE S1:** Number of individuals of *Gobio occitaniae*, *Barbatula barbatula*, *Squalius cephalus*, *Phoxinus phoxinus*, *Leuciscus burdigalensis* and *Parachondrostoma toxostoma* genotyped *per* sampling site, along with sampling sites information. N is the number of sampling sites successfully sampled for genetic analyses. Sites successfully sampled for genetic analyses are sites where at least ten individuals (or six individuals for the two rare species *L. burdigalensis* and *Pa. toxostoma*) have been successfully genotyped.

| River | Location | Year | Longitude | Latitude | Number of individuals genotyped | | | | | |
|---|---|---|---|---|---|---|---|---|---|---|
| | | | | | *Gobio occitaniae N=83* | *Barbatula barbatula N=48* | *Squalius cephalus N=66* | *Phoxinus phoxinus N=63* | *Leuciscus burdigalensis N=29* | *Parachondrostoma toxostoma N=14* |
| Agoût | Bez | 2010 | N 43°37'28.1" | E 002°29'55.2" | 13 | — | — | 24 | — | — |
| Agoût | Lavaur | 2011 | N 43°42'33.3" | E 001°47'20.8" | 25 | — | 25 | — | — | — |
| Agoût | La Salvetat-sur-Agoût | 2011 | N 43°36'22.3" | E 002°42'03.0" | 11 | — | — | 25 | — | — |
| Ariège | Tarrascon-sur-Ariège | 2011 | N 42°50'48.9" | E 001°36'19.3" | — | 10 | — | 25 | — | — |
| Ariège | Venerque | 2010 | N 43°26'13.8" | E 001°26'15.5" | 25 | 25 | — | 30 | — | — |
| Ariège | Le Vernet | 2011 | N 43°11'12.0" | E 001°35'59.8" | 25 | 25 | — | 25 | — | — |
| Arize | Mas-d'Azil | 2011 | N 43°05'03.8" | E 001°22'25.3" | 25 | 25 | — | 25 | 6 | — |
| Auvézère | Génis | 2011 | N 45°20'38.2" | E 001°10'25.9" | 25 | — | 25 | — | — | — |
| Avance | Casteljaloux | 2010 | N 44°18'54.7" | E 000°05'28.8" | — | — | — | — | — | — |
| Avance | Montpouillan | 2010 | N 44°28'04.2" | E 000°07'33.1" | — | 25 | 16 | 25 | — | — |
| Aveyron | Druelle | 2010 | N 44°20'12.4" | E 002°29'29.2" | 24 | 19 | 25 | 25 | 10 | — |
| Aveyron | Féneyrols | 2010 | N 44°07'52.0" | E 001°48'51.0" | 25 | — | 25 | 25 | 7 | 6 |
| Aveyron | Gaillac-d'Aveyron | 2010 | N 44°21'27.8" | E 002°54'27.8" | 25 | 25 | 25 | 25 | — | — |
| Aveyron | Monteils | 2010 | N 44°17'09.1" | E 002°00'07.5" | 25 | 25 | 25 | 25 | — | — |
| Grande Baïse | Brouilh-Monbert | 2011 | N 43°40'14.0" | E 000°23'56.0" | 15 | — | 25 | — | — | — |
| Grande Baïse | Nérac | 2011 | N 44°08'50.7" | E 000°20'09.2" | — | — | 25 | — | — | — |
| Barguelonne | Montbarla | 2011 | N 44°12'35.0" | E 001°03'40.6" | 25 | 16 | 25 | 25 | — | 9 |
| Célé | Amis du Célé | 2010 | N 44°31'13.0" | E 001°40'42.6" | 25 | 25 | 16 | 25 | 14 | 25 |
| Célé | Bagnac | 2010 | N 44°39'51.9" | E 002°10'05.6" | 25 | 15 | — | 25 | 10 | — |
| Célé | Boussac | 2010 | N 44°35'46.2" | E 001°55'02.4" | 24 | 10 | 24 | 25 | 25 | 7 |
| Célé | Moulin de Claude upstream | 2011 | N 44°41'00.5" | E 002°16'37.4" | 28 | — | — | — | — | — |
| Célé | Moulin de Claude downstream | 2011 | N 44°41'00.5" | E 002°16'37.4" | 30 | — | — | — | — | — |

| | | | | | | | | | | |
|---|---|---|---|---|---|---|---|---|---|---|
| Célé | Sainte-Eulalie | 2010 | N 44°35'36.6" | E 001°52'25.7" | 25 | — | 11 | 25 | 15 | 8 |
| Célé | Merlie | 2010 | N 44°31'59.4" | E 001°44'48.6" | 25 | 25 | 16 | 24 | 25 | — |
| Cère | Bretenoux | 2010 | N 44°54'57.5" | E 001°50'14.9" | 25 | 25 | 10 | 25 | 9 | — |
| Ciron | Préchac | 2011 | N 44°24'35.5" | W 000°20'25.0" | 23 | 16 | — | 25 | — | — |
| Corrèze | Les Angles-sur-Corrèze | 2011 | N 45°18'06.2" | E 001°47'49.6" | 25 | — | — | 25 | — | — |
| Dadou | Montdragon | 2011 | N 43°46'40.7" | E 002°05'47.1" | 25 | — | 25 | 25 | 25 | — |
| Dadou | Realmont | 2010 | N 43°46'45.2" | E 002°10'54.3" | 22 | — | 20 | — | 19 | — |
| Dordogne | Brivezac | 2011 | N 45°01'25.4" | E 001°50'36.6" | 25 | 25 | — | 25 | — | — |
| Dordogne | Cénac | 2011 | N 44°48'29.1" | E 001°11'32.2" | 25 | 25 | 25 | 25 | 26 | — |
| Dordogne | Chalameyroux | 2011 | N 45°35'08.9" | E 002°32'57.8" | — | — | — | 19 | — | — |
| Dordogne | Cours-de-pile | 2011 | N 44°51'10.4" | E 000°33'33.0" | 25 | — | 25 | — | — | — |
| Dordogne | Fleix | 2011 | N 44°51'44.9" | E 000°14'35.7" | 25 | — | 25 | — | — | — |
| Dordogne | Meyraguet | 2010 | N 44°51'08.6" | E 001°32'38.6" | 23 | 25 | — | 25 | — | — |
| Dordogne | Prudhomat | 2010 | N 44°53'49.5" | E 001°47'20.2" | 25 | 25 | 16 | 25 | 14 | — |
| Dourdou | Montlaur | 2010 | N 43°53'02.4" | E 002°50'19.8" | 25 | 24 | 25 | 25 | 9 | — |
| Dropt | Cavarc | 2010 | N 44°39'32.7" | E 000°38'53.4" | 25 | 20 | 25 | — | — | — |
| Dronne | Coutras | 2011 | N 45°02'24.9" | W 000°07'52.4" | 25 | — | 24 | — | — | — |
| Dronne | Saint-Front-la-Riviere | 2011 | N 45°28'09.3" | E 000°43'28.5" | 25 | 25 | 25 | 24 | 15 | — |
| Dronne | Valeuil | 2011 | N 45°20'05.4" | E 000°36'50.6" | 25 | 25 | 25 | 25 | 16 | — |
| Eau blanche | Moulin Noir | 2011 | N 44°45'24.7" | W 000°32'25.0" | 25 | 10 | 25 | — | 7 | — |
| Elle | Terrasson-Lavilledieu | 2011 | N 45°08'51.4" | E 001°15'37.6" | 25 | 25 | 25 | 25 | 15 | 25 |
| Garonne | Agen | 2010 | N 44°12'25.7" | E 000°36'21.5" | 25 | — | 26 | — | — | — |
| Garonne | Bourret | 2010 | N 43°57'16.2" | E 001°10'20.4" | 25 | — | 25 | — | — | — |
| Garonne | Clarac | 2011 | N 43°05'52.3" | E 000°37'45.3" | 20 | 25 | — | 25 | — | — |
| Garonne | Couthures-sur-Garonne | 2010 | N 44°30'52.3" | E 000°04'49.1" | 25 | — | 25 | — | — | — |
| Garonne | Gagnac-sur-Garonne | 2010 | N 43°42'18.1" | E 001°21'52.8" | 25 | — | 25 | 15 | — | — |
| Garonne | Saint-Julien-sur-Garonne | 2010 | N 43°14'34.7" | E 001°09'19.0" | 25 | 24 | — | 25 | — | — |
| Garonne | Muret | 2010 | N 43°27'36.5" | E 001°19'52.7" | 25 | — | 25 | 25 | — | — |
| Gers | Aurenque | 2011 | N 43°53'19.7" | E 000°39'29.2" | 24 | — | 25 | — | — | — |
| Gers | Chélan | 2011 | N 43°20'32.5" | E 000°33'00.6" | 14 | 20 | — | 25 | — | — |
| Gers | Preignan | 2011 | N 43°43'02.2" | E 000°37'18.8" | 25 | 25 | 18 | — | — | — |

| | | | | | | | | | | |
|---|---|---|---|---|---|---|---|---|---|---|
| Grand Hers | Besset | 2010 | N 43°05'03.2" | E 001°50'24.2" | 25 | 25 | 16 | 25 | — | — |
| Grand Hers | Calmont | 2010 | N 43°17'10.2" | E 001°37'59.1" | 25 | — | 25 | 25 | — | 25 |
| Isle | Saint-Laurent-des-Hommes | 2011 | N 45°01'46.3" | E 000°14'35.2" | — | — | 25 | — | — | — |
| Isle | Trélissac | 2011 | N 45°11'56.7" | E 000°48'04.8" | 25 | 25 | 25 | 25 | 25 | — |
| Lot | Balsièges | 2011 | N 44°28'43.9" | E 003°27'09.0" | 25 | 25 | 14 | 25 | — | — |
| Lot | Bouziès | 2010 | N 44°29'07.6" | E 001°37'44.4" | 24 | — | 23 | 18 | 10 | — |
| Lot | Labéraudie | 2010 | N 44°26'27.6" | E 001°26'23.8" | 25 | — | 25 | — | — | — |
| Lot | Clairac | 2011 | N 44°20'50.1" | E 000°22'10.8" | 16 | — | 25 | — | — | — |
| Lot | Entraygues-sur-Truyère | 2010 | N 44°37'38.1" | E 002°34'40.4" | 25 | 14 | 25 | 25 | — | — |
| Lot | Pomeyrols | 2010 | N 44°28'11.0" | E 003°01'17.3" | 25 | 20 | — | 25 | — | — |
| Lot | Livinhac-le-Haut | 2010 | N 44°35'06.6" | E 002°13'50.6" | 25 | 23 | 25 | 25 | — | — |
| Louge | Fousseret | 2010 | N 43°16'59.9" | E 001°04'49.4" | 24 | 24 | — | 25 | — | 8 |
| Osse | Monclar-sur-Losse | 2011 | N 43°31'51.1" | E 000°20'08.4" | 25 | 25 | 25 | — | — | — |
| Osse | Mouchan | 2011 | N 43°54'21.0" | E 000°17'36.4" | 25 | — | 25 | 25 | — | — |
| Salat | Saint-Girons | 2011 | N 42°59'07.7" | E 001°08'33.3" | 20 | — | — | 25 | — | — |
| Salat | Touille | 2010 | N 43°04'38.7" | E 000°58'05.6" | 24 | 25 | 23 | 25 | 14 | 25 |
| Save | Espaon | 2011 | N 43°25'20.1" | E 000°51'21.6" | 25 | — | 25 | 21 | 6 | 18 |
| Save | Lévignac | 2010 | N 43°40'05.0" | E 001°11'25.4" | 25 | — | 25 | — | — | — |
| Ségur | Les Bardes | 2011 | N 44°38'03.0" | W 000°02'01.5" | 15 | — | — | — | — | — |
| Ségur | Moulin de Mardon | 2011 | N 44°39'33.1" | W 000°01'38.5" | 24 | 15 | — | 19 | — | — |
| Tarn | Avalats | 2011 | N 43°56'32.7" | E 002°14'26.7" | 25 | 25 | 24 | 21 | — | — |
| Tarn | Brousse-le-Château | 2010 | N 44°00'30.6" | E 002°36'28.6" | 25 | — | 25 | 25 | — | — |
| Tarn | Formarès | 2011 | N 44°20'18.6" | E 003°34'59.9" | 25 | 25 | 25 | 25 | 13 | — |
| Tarn | Millau | 2010 | N 44°06'29.7" | E 003°05'08.6" | 25 | 25 | 25 | 25 | — | — |
| Tarn | Moissac | 2010 | N 44°05'52.4" | E 001°05'10.1" | — | — | 25 | — | — | — |
| Tarn | Rabastens | 2010 | N 43°49'04.9" | E 001°43'29.5" | 25 | — | 25 | — | — | — |
| Tarn | Villemur-sur-Tarn | 2011 | N 43°52'31.2" | E 001°29'51.4" | 25 | — | 25 | — | — | — |
| Touch | Lamasquère | 2010 | N 43°29'32.1" | E 001°13'48.5" | 24 | 16 | 25 | 23 | 8 | — |
| Truyère | Loubaresse | 2011 | N 44°56'18.4" | E 003°13'59.6" | 16 | — | 26 | — | — | — |
| Truyère | Valcaylès | 2011 | N 44°45'38.3" | E 002°40'49.5" | — | — | — | 25 | — | — |
| Vère | Cahuzac-sur-Vère | 2010 | N 43°58'39.6" | E 001°54'51.0" | 25 | 25 | 25 | 25 | — | — |

| | | | | | | | | | | |
|---|---|---|---|---|---|---|---|---|---|---|
| Vézère | Saint-Léon-sur-Vézère | 2011 | N 45°00'36.8" | E 001°06'31.7" | 24 | — | 25 | — | — | — |
| Vézère | Treignac | 2011 | N 45°32'20.6" | E 001°47'38.5" | — | — | — | 25 | — | — |
| Viaur | Bannès | 2010 | N 44°15'57.7" | E 002°39'05.7" | 25 | 14 | — | 25 | 8 | — |
| Viaur | Calquière | 2010 | N 44°09'12.3" | E 002°12'15.9" | 25 | — | 25 | 25 | 14 | 13 |
| Viaur | Capelle | 2010 | N 44°15'23.7" | E 002°37'02.8" | 25 | 25 | 25 | 25 | 19 | — |
| Viaur | Saint-Just | 2010 | N 44°07'24.6" | E 002°21'57.0" | 25 | 12 | 25 | 25 | 17 | 23 |
| Viaur | Navech | 2010 | N 44°09'25.5" | E 002°23'18.3" | 25 | 18 | 25 | 25 | 12 | 25 |
| Viaur | Serres | 2010 | N 44°12'29.4" | E 002°31'25.6" | 25 | — | 25 | 25 | — | 6 |

**TABLE S2:** Mean values of Allelic Richness (*AR*), Private Allelic Richness (*PA*) and $D_{est}$ calculated across sampling sites for the six species.

| | Genetic index | | |
|---|---|---|---|
| Species name | *AR* | *PA* | $D_{est}$ |
| *Squalius cephalus* | 3.829 | 0.059 | 0.164 |
| *Gobio occitaniae* | 5.268 | 0.056 | 0.200 |
| *Barbatula barbatula* | 4.441 | 0.099 | 0.383 |
| *Phoxinus phoxinus* | 5.821 | 0.147 | 0.267 |
| *Leuciscus burdigalensis* | 3.470 | 0.162 | 0.169 |
| *Parachondrostoma toxostoma* | 2.114 | 0.036 | 0.069 |

**TABLE S3:** Pearson's correlation coefficients between values of *AR*, *PA* and $D_{est}$ observed at the sampling site level for one common species, and those observed for other common species. Bold correlation values are significant. * p-value <0.05; ** p-value <0.01; *** p-value <0.001.

| Genetic index | Species name | Species name | | | |
|---|---|---|---|---|---|
| | | *Squalius cephalus* | *Gobio occitaniae* | *Barbatula barbatula* | *Phoxinus phoxinus* |
| *AR* | *Squalius cephalus* | — | | | |
| | *Gobio occitaniae* | 0.041 | — | | |
| | *Barbatula barbatula* | -0.047 | **0.667*** ** | — | |
| | *Phoxinus phoxinus* | 0.277 | **0.559*** ** | **0.718*** ** | — |
| *PA* | *Squalius cephalus* | — | | | |
| | *Gobio occitaniae* | -0.042 | — | | |
| | *Barbatula barbatula* | 0.189 | 0.040 | — | |
| | *Phoxinus phoxinus* | -0.122 | -0.068 | 0.023 | — |
| $D_{est}$ | *Squalius cephalus* | — | | | |
| | *Gobio occitaniae* | 0.071 | — | | |
| | *Barbatula barbatula* | **0.352*** | **0.403**** | — | |
| | *Phoxinus phoxinus* | 0.078 | **0.485*** ** | **0.347*** | — |

**TABLE S4:** Table reporting for each common species the number of sampling sites (and the proportion of the total number of sampled sites) that showed an irreplaceability value of 100% when minimum sets of sampling sites representing (i) 50 %, (ii) 75%, (iii) 90% or (iv) 100% of the total number of alleles present in the river basin were identified with the software Marxan. The selection of a site means that the site is completely irreplaceable for attaining the aimed conservation target (i.e. 50, 75, 90 or 100% of alleles).

| Species name<br>Number of total sampled sites | *Squalius cephalus*<br>n=66 | *Gobio occitaniae*<br>n=83 | *Barbatula barbatula*<br>n=48 | *Phoxinus phoxinus*<br>n=63 |
|---|---|---|---|---|
| Conservation target of 50% of alleles | 1 (1.52%) | 0 | 0 | 1 (1.59%) |
| Conservation target of 75% of alleles | 2 (3.03%) | 1 (1.20%) | 0 | 4 (6.35%) |
| Conservation target of 90% of alleles | 7 (10.61%) | 3 (3.61%) | 9 (18.85%) | 18 (28.57%) |
| Conservation target of 100% of alleles | 26 (39.39%) | 21 (25.30%) | 20 (41.67%) | 43 (68.26%) |

**TABLE S5:** Results of Generalized Linear Models (GLMs) aimed at testing whether the positional importance of sampling sites across the riverscape (i.e. measured as their betweenness centrality values) and the distance of the sampling site to the outlet may predict irreplaceable sites at the 90% conservation target. Results are presented for each of the six species independently. Estimates are parameter values indicating the sign (and strength) of the relationships between the response variable (the probability for a site to be a priority site) and each explanatory variable. Z values correspond to the statistic used to test the significance of each relationship. Bold p-values are significant.

| Species | Explanatory variables | Estimates | Z values | p-values |
|---|---|---|---|---|
| *(a) Squalius cephalus* | *Distance* | -1.912e-06 | -0.609 | 0.543 |
| | *Betweenness* | -3.430e+03 | -1.210 | 0.226 |
| *(b) Gobio occitaniae* | *Distance* | 2.994e-06 | 0.585 | 0.558 |
| | *Betweenness* | -3.674e+02 | -0.156 | 0.876 |
| *(c) Barbatula barbatula* | *Distance* | -3.483e-06 | -1.233 | 0.218 |
| | *Betweenness* | -2.628e+03 | -1.277 | 0.202 |
| *(d) Phoxinus phoxinus* | ***Distance*** | **6.856e-06** | **2.572** | **0.010*** |
| | *Betweenness* | 1.327e+03 | 1.381 | 0.167 |
| *(e) Leuciscus burdigalensis* | *Distance* | -1.806e-06 | -0.702 | 0.483 |
| | *Betweenness* | -4.770e+02 | -0.386 | 0.700 |
| *(f) Parachondrostoma toxostoma* | *Distance* | -3.399e-06 | -0.747 | 0.455 |
| | *Betweenness* | -4.472e+03 | -0.851 | 0.395 |

**TABLE S6:** Table reporting for each species the percentages of their total number of alleles that are secured by the sampling sites having been identified as irreplaceable sites for the other species (considering the 90% conservation target).

| Proportion of alleles from (below) that fell within irreplaceable sites from (right): | *Squalius cephalus n=7* | *Gobio occitaniae n=3* | *Barbatula barbatula n=9* | *Phoxinus phoxinus n=18* | *Leuciscus burdigalensis n=11* | *Parachondrostoma toxostoma n=3* | Common species *n=32* | Rare species *n=14* |
|---|---|---|---|---|---|---|---|---|
| *Squalius cephalus* | *90.00* | 17.66 | 61.52 | 56.52 | 61.60 | 39.73 | — | 65.18 |
| *Gobio occitaniae* | 65.60 | *90.00* | 66.76 | 79.58 | 70.27 | 50.97 | — | 73.40 |
| *Barbatula barbatula* | 49.79 | 31.65 | *90.00* | 79.57 | 76.78 | 38.95 | — | 79.40 |
| *Phoxinus phoxinus* | 48.25 | 42.60 | 68.43 | *90.00* | 63.49 | 44.61 | — | 68.88 |
| *Leuciscus burdigalensis* | 44.73 | 0 | 66.13 | 67.09 | *90.00* | 36.42 | 81.79 | — |
| *Parachondrostoma toxostoma* | 78.55 | 0 | 78.55 | 50.00 | 67.14 | *90.00* | 90.00 | — |
| *Average over all other species* | *55.58* | *18.38* | *68.28* | *66.55* | *67.86* | *42.14* | | |

**TABLE S7:** Results of Generalized Linear Models (GLMs) aimed at testing which of genetic indices among *AR*, *PA* and $D_{est}$ better predict irreplaceable sites at the 90% conservation target. Results are presented for each of the six species independently. Estimates are parameter values indicating the sign (and strength) of the relationships between the response variable (the probability for a site to be a priority site) and each explanatory variable. Z values correspond to the statistic used to test the significance of each relationship. Bold p-values are significant.

| Species | Explanatory variables | Estimates | Z values | p-values |
|---|---|---|---|---|
| *(a) Squalius cephalus* | *AR* | 0.445 | 0.306 | 0.759 |
| | ***PA*** | **6.115** | **2.041** | **0.041*** |
| | $D_{est}$ | 0.007 | 0.005 | 0.996 |
| *(b) Gobio occitaniae* | *AR* | -7.318 | -0.799 | 0.424 |
| | *PA* | 6.514 | 1.255 | 0.209 |
| | $D_{est}$ | -10.165 | -0.836 | 0.403 |
| *(c) Barbatula barbatula* | *AR* | -0.069 | -0.092 | 0.926 |
| | ***PA*** | **1.644** | **2.630** | **0.009**** |
| | $D_{est}$ | -0.741 | -0.714 | 0.475 |
| *(d) Phoxinus phoxinus* | *AR* | 0.119 | 0.210 | 0.834 |
| | ***PA*** | **3.123** | **3.834** | **<0.001***** |
| | $D_{est}$ | -0.851 | -1.396 | 0.163 |
| *(e) Leuciscus burdigalensis* | *AR* | 1.868 | 1.004 | 0.315 |
| | ***PA*** | **3.383** | **2.249** | **0.025*** |
| | $D_{est}$ | 0.969 | 0.659 | 0.510 |
| *(f) Parachondrostoma toxostoma* | *AR* | -1.977 | -1.013 | 0.311 |
| | *PA* | 4.975 | 1.330 | 0.184 |
| | $D_{est}$ | -1.341 | -0.806 | 0.420 |

**APPENDIX A1:** Detailed information on the genotyping procedure.

Genomic DNA was extracted using a salt-extraction protocol (Aljanabi and Martinez 1999). We used multiplexed Polymerase Chain Reactions (PCRs) to co-amplify 8-15 microsatellite loci depending on the species (8 for *G. occitaniae*, 9 for *B. barbatula*, 10 for *S. cephalus* and *Ph. phoxinus*, 14 for *L. burdigalensis* and 15 for *Pa. toxostoma*). We used 5-20 ng of genomic DNA and QIAGEN® Multiplex PCR Kits (Qiagen, Valencia, CA, USA) to perform PCR amplifications. Details on loci, primer concentrations, and multiplex sets are given in the table below. The genotyping was conducted on an ABI PRISM™ 3730 Automated Capillary Sequencer (Applied Biosystems, Foster City, CA, USA), and the scoring of allele sizes was done using GENEMAPPER® v.4.0 (Applied Biosystems).

| Species | Locus | GenBank Accession ID | Fluorescent Dye | Multiplex Kit | Primer Concentration (nM) | Reference |
|---|---|---|---|---|---|---|
| *Squalius cephalus* | Ca01 | AF277573 | HEX | 1 | 100 | Dimsoski et al. 2000 |
| | CypG30 | AY439148 | NED | 1 | 150 | Baerwald et al. 2004 |
| | Lc293 | EF362795 | FAM | 1 | 100 | Vyskocilova et al. 2007 |
| | Lcel100 | AY962249 | FAM | 1 | 300 | Larno et al. 2005 |
| | Lc290 | EF362794 | HEX | 1 | 100 | Vyskocilova et al. 2007 |
| | CypG24 | AY439142 | FAM | 2 | 50 | Baerwald et al. 2004 |
| | CypG27 | AY439145 | HEX | 2 | 200 | Baerwald et al. 2004 |
| | Lc27 | EF362792 | NED | 2 | 100 | Vyskocilova et al. 2007 |
| | LceC1 | AY962241 | FAM | 2 | 100 | Larno et al. 2005 |
| | Lid11 | AB112736 | FAM | 2 | 250 | Barinova et al. 2004 |
| *Gobio occitaniae* | Ca01 | AF277573 | HEX | 1 | 100 | Dimsoski et al. 2000 |
| | Gob16 | DQ207803 | HEX | 1 | 30 | Knapen et al. 2006 |
| | Rhca20 | DQ106915 | NED | 1 | 100 | Girard and Angers 2006 |
| | Gob12 | DQ207801 | FAM | 1 | 175 | Knapen et al. 2006 |
| | Gob15 | DQ207802 | HEX | 2 | 100 | Knapen et al. 2006 |
| | Gob22 | DQ207804 | FAM | 2 | 50 | Knapen et al. 2006 |
| | Gob28 | DQ207805 | HEX | 2 | 100 | Knapen et al. 2006 |
| | MFW1 | N.A. | NED | 2 | 100 | Crooijmans et al. 1997 |
| *Barbatula barbatula* | Bbar1 | AF311346 | NED | 1 | 40 | Taylor et al. 2001 |
| | Bbar7 | AF310881 | FAM | 1 | 80 | Taylor et al. 2001 |
| | Bbar8 | AF310882 | HEX | 1 | 200 | Taylor et al. 2001 |
| | Lec12 | AB286043 | NED | 1 | 80 | Koizumi et al. 2007 |
| | Bbar11 | AF310883 | HEX | 2 | 80 | Taylor et al. 2001 |
| | Bbar9 | AF311349 | NED | 2 | 60 | Taylor et al. 2001 |
| | Lec01 | AB286032 | FAM | 2 | 80 | Koizumi et al. 2007 |
| | Lec05 | AB286036 | HEX | 2 | 150 | Koizumi et al. 2007 |
| *Phoxinus phoxinus* | CypG27 | AY439145 | HEX | 1 | 200 | Baerwald et al. 2004 |
| | CypG30 | AY439148 | NED | 1 | 100 | Baerwald et al. 2004 |
| | LceC1 | AY962241 | FAM | 1 | 50 | Larno et al. 2005 |
| | Lid2 | AB112733 | FAM | 1 | 200 | Barinova et al. 2004 |
| | Rru4 | AB112740 | HEX | 1 | 20 | Barinova et al. 2004 |
| | Ca12 | AF277584 | FAM | 2 | 130 | Dimsoski et al. 2000 |
| | CypG03 | AY439122 | HEX | 2 | 50 | Baerwald et al. 2004 |
| | Lc27 | EF362792 | NED | 2 | 100 | Vyskocilova et al. 2007 |
| | MFW1 | N.A. | HEX | 2 | 250 | Crooijmans et al. 1997 |
| | Rhca20 | DQ106915 | FAM | 2 | 50 | Girard and Angers 2006 |
| *Leuciscus burdigalensis* | Lid8 | AB112735 | FAM | 1 | 100 | Barinova et al. 2004 |
| | Lc290 | EF362794 | HEX | 1 | 100 | Vyskocilova et al. 2007 |
| | CypG30 | AY439148 | NED | 1 | 200 | Baerwald et al. 2004 |
| | LceC1 | AY962241 | FAM | 2 | 50 | Larno et al. 2005 |
| | Lid2 | AB112733 | FAM | 2 | 80 | Barinova et al. 2004 |
| | Lid11 | AB112736 | FAM | 2 | 150 | Barinova et al. 2004 |
| | CypG03 | AY439122 | HEX | 2 | 50 | Baerwald et al. 2004 |
| | Lc27 | EF362792 | NED | 4 | 150 | Vyskocilova et al. 2007 |
| | Ca12 | AF277584 | FAM | 3 | 150 | Dimsoski et al. 2000 |
| | Rru4 | AB112740 | HEX | 3 | 50 | Barinova et al. 2004 |

| | | | | | | |
|---|---|---|---|---|---|---|
| | Z21908 | G40277 | NED | 3 | 100 | Shimoda et al. 1999 |
| | Lco5 | AY318781 | FAM | 4 | 150 | Turner et al. 2004 |
| | MFW1 | N.A. | HEX | 4 | 200 | Crooijmans et al. 1997 |
| | Rhca20 | DQ106915 | FAM | 4 | 150 | Girard and Angers 2006 |
| *Parachondrostoma toxostoma* | BL1-30 | FJ468353 | 6-FAM | 1 | 100 | Dubut et al. 2009a |
| | Rser10 | AJ312850 | 6-FAM | 1 | 100 | Dawson et al. 2003 |
| | LleC-090 | FJ601722 | 6-FAM | 1 | 100 | Dubut et al. 2009b |
| | LceC1 | AY962241 | HEX | 1 | 100 | Larno et al. 2005 |
| | BL1-61 | FJ468351 | HEX | 1 | 75 | Dubut et al. 2009a |
| | Ca01 | AF277573 | ATTO550 | 1 | 300 | Dimsoski et al. 2000 |
| | N7K4 | AJ566138 | ATTO550 | 1 | 100 | Mesquita et al. 2003 |
| | CtoA-256 | GU254032 | 6-FAM | 2 | 150 | Dubut et al. 2010 |
| | BL1-2b | FJ468347 | 6-FAM | 2 | 100 | Dubut et al. 2009a |
| | Rru4 | AB112740 | 6-FAM | 2 | 100 | Barinova et al. 2004 |
| | Lsou34 | EF209012 | HEX | 2 | 75 | Muenzel et al. 2007 |
| | LleA029 | FJ601714 | HEX | 2 | 100 | Dubut et al. 2009b |
| | CtoF172 | GU254034 | ATTO550 | 2 | 100 | Dubut et al. 2010 |
| | Lsou05 | EF209002 | ATTO550 | 2 | 100 | Muenzel et al. 2007 |
| | BL1-T2 | FJ468348 | ATTO550 | 2 | 100 | Dubut et al. 2009a |

N.A. means not available.

**APPENDIX A2:** Detailed information on the genotyping quality control procedures.

We determined for each of the six species the occurrence of null alleles and potential scoring errors with the program MICROCHECKER 2.3 (Van Oosterhout et al. 2004). We tested for departures from Hardy-Weinberg (HW) equilibrium with the 'adegenet' R package v1.6-2 (Jombart 2008). The program GENEPOP v4.0 (Rousset 2008) was used to assess linkage disequilibrium among loci within sites. Levels of significance for these multiple tests were adjusted using Bonferroni procedures. We found significant deviations from HW after Bonferroni corrections for a few locus/population pairs for each species (i.e. 15/378 significant pairs for *Barbatula barbatula*; 7/592 for *Gobio occitaniae;* 19/540 for *Phoxinus phoxinus,* 24/600 for *Squalius cephalus,* 3/406 for *Leuciscus burdigalensis,* and 4/210 for *Parachondrostoma toxostoma*). However, these departures were not consistently clustered among loci or populations for any species (see also Supplementary Data File 1 for raw results). We also detected a few significant linkage disequilibrium between pairs of loci after Bonferroni corrections for some populations for each species excepting *Pa. toxostoma* and *L. burdigalensis* (i.e. 4/1512 significant tests for *B. barbatula*; 6/2072 for *G. occitaniae*; 5/2430 for *Ph. phoxinus* and 6/2700 for *S. cephalus*), but no clear patterns of linkage disequilibrium appeared for any species across populations (Supplementary Data File 1). We did not find evidence for scoring errors due to stuttering or large allele dropouts in our datasets, although Microchecker indicated some significant homozygote excesses in a few cases (i.e. 7/378 significant homozygote excesses for *B. barbatula*; 5/592 for *G. occitaniae*; 20/540 for *Ph. phoxinus* and 7/600 for *S. cephalus*). However, these homozygote excesses were not consistent for any loci across populations within each species. Given the small extent of these deviations and the large spatial extent of the databases, we assumed that they weakly affected our main findings.

**APPENDIX A3:** Detailed information on the building of GLMSSNs

We used geostatistical modelling tools to explore spatial patterns of $\alpha$ and $\beta$ intraspecific genetic diversity for the four common species at the riverscape scale by predicting the distributions of *AR*, *PA* and $D_{est}$ from the observed empirical values using Generalized Linear Models for Spatial Stream Networks (GLMSSN; Peterson & Ver Hoef 2014; Ver Hoef et al. 2014). This was done using the 'STARS' toolset of ARCGIS v10.2 and the R package 'SSN' (Peterson & Ver Hoef 2014; Ver Hoef et al. 2014). We conducted a model selection procedure based on a comparison of Akaike Information Criteria (AIC) estimated for several competing GLMSSNs. These models were built by (i) assuming three geographic descriptors (i.e. topological distance from the outlet, longitude and latitude) as explanatory variables, and (ii) choosing a tail-down covariance structure model among the following ones: exponential, Mariah, spherical, linear-with-sill and Epanechnikov. The model with the lowest AIC score (see Supplementary Data File 1 for raw results) was considered as the best model and was used to estimate the slope and the significance of the relationships between explanatory variables and each genetic index, which allowed testing whether or not spatial patterns of intraspecific genetic diversity can be detected. We finally used predictions from the best models to produce krigged maps for each common species and each genetic index, to visually represent the spatial distribution of intraspecific genetic diversity across the whole river drainage, and to visually highlight hot- and cold-spots of intraspecific genetic diversity.

**APPENDIX A4:** Detailed information on the SCP procedures

We tested whether conservation targets (i.e. the percentage of total number of conservation units to be present in the final conservation solution) and analytical approaches (i.e. species-specific or species-pooled analyses) influence the identification of irreplaceable sites. Systematic conservation planning (SCP) methods traditionally use species presence/absence data as input data to identify irreplaceable sites for the conservation of taxonomic diversity at the community level (i.e. sites that cannot be excluded from an optimal selection of sites for conservation; Ball et al. 2009). Here, we replaced species presence/absence data with allele presence-absence data as in Diniz-Filho et al. (2012) to identify irreplaceable sites for the conservation of intraspecific genetic diversity of each species in the river drainage. We used the program Marxan v2.1 (Ball et al. 2009) and genotypic data from the common species to identify, for each common species independently, an optimal set of sites that best represent at least 50, 75, 90 or 100% of the total number of alleles present in the whole riverscape at a minimum "cost", which corresponds to four conservation targets. Given the lack of ground estimates for conservation cost, we used a constant cost *per* site, so our objective translated into identifying the minimum number of sites that represent a given proportion of intraspecific genetic diversity (i.e. 50, 75, 90 or 100% of the total pool of alleles represented at least once; Hermoso & Kennard 2012). We arbitrarily choose the 50, 75 and 100% conservation targets to explore how the proportion of alleles to protect affects the selection of irreplaceable sites. We additionally tested the 90% conservation target, as it corresponds to a threshold target being typically assumed in *ex-situ* conservation plans (Frankham et al. 2002). In order to estimate the relevance of each site to preserve a given proportion of the allelic diversity in the river basin, we used two different methods. In case of 100% of allelic diversity, we used the traditional irreplaceability measure reported by Marxan that ranges between 0% (not irreplaceable) and 100% (highly irreplaceable). This measure is estimated by running the optimization algorithm a number of times (N=100 runs in our case) and then computing the frequency of selection of each site within the solutions obtained. Sites with unique allelic composition will be selected across all runs and reported as highly irreplaceable, whereas sites with more common alleles, replaceable by other sites with the same alleles, will appear poorly irreplaceable. For the other conservation targets, we selected a random pull of 50, 75 and 90% of the total number of alleles existing at the basin level and for each species. We then ran the optimization algorithm to identify the minimum set of sites representing this particular selection of alleles. Given that the analyses were run only on a subset of alleles, we replicated this process 100 times to minimize the effect of the arbitrary selection of alleles. For each subset of alleles, we ran the Marxan procedure as explained above (e.g. constant cost and 100 runs for each) and retained the best solution for subsequent analyses (solution with the lowest value for the objective function). We then calculated the irreplaceability as the frequency of selection of each site within the 100 random pull of alleles. For a given species and a given conservation target, we considered a site as irreplaceable for genetic diversity conservation when it displayed an irreplaceability value of 100%. We selected such a high threshold so as to be conservative. We tested and compared visually how the proportions of irreplaceable sites vary among species and conservation targets.
